# Mutational insights among the structural proteins of SARS-CoV-2: frequencies and evolutionary trends in American countries

**DOI:** 10.1101/2022.06.22.497134

**Authors:** Mohammad Abavisani, Karim Rahimian, Reza Khayami, Mahsa Mollapour Sisakht, Mohammadamin Mahmanzar, Zahra Meshkat

## Abstract

Severe acute respiratory syndrome coronavirus 2 (SARS-CoV-2) has a role in the mortality of more than 6 million people worldwide. This virus owns the genome, which contains four structural proteins, including spike (S), envelope (E), membrane (M), and nucleocapsid (N). The occurrence of structural mutations can induce the emergence of new variants. Depending on the mutations, the variants may display different patterns of infectivity, mortality, and sensitivity toward drugs and vaccines. In this study, we analyzed samples of amino-acid sequences (AASs) for structural proteins from the coronavirus 2019 (COVID-19) declaration as a pandemic to April 2022 among American countries. The analysis process included considering mutations’ frequencies, locations, and evolutionary trends utilizing sequence alignment to the reference sequence. In the following, the results were compared with the same analyses among the samples of the entire world. Results displayed that despite samples of North America and international countries that own the region of 508 to 635 with the highest mutation frequency among S AASs, the region with the same characteristic was concluded as 1 to 127 in South America. Besides, the most frequent mutations in S, E, M, and N proteins from North America and worldwide samples were concluded as D614G, T9I, I82T, and R203M. In comparison, R203K was the first frequent mutation in N samples in South America. Widely comparing mutations between North America and South America and between the Americas and the world can help scientists introduce better drug and vaccine development strategies.

## Introduction

Severe acute respiratory syndrome coronavirus 2 (SARS-CoV-2) is a member of the *Betacoronavirus* genus of the *Coronaviridae* family. Viruses of this family have positive-sense single-stranded enveloped RNA genomes, ranging from 26 to 32 kilobases in length [1]. Since the outbreak started in Wuhan, China, more than 6 million people have been killed by the Coronavirus 2019 (COVID-19) pandemic worldwide [2].

To date, 14 open reading frames (ORFs) encoding 27 structural and non-structural proteins (NSPs) have been identified for SARS-CoV-2. The 5’ end of the SARS-CoV-2 genome contains genes that encode pp1ab and pp1a proteins that together include 15 NSPs (NSP1-NSP10 and NSP12-NSP16), while the 3’ end of its genome contains genes that encode four structural proteins called spike (S), envelope (E), membrane (M) and nucleocapsid (N) as well as eight accessory proteins called 3a, 3b, p6, 7a, 7b, 8b, 9b, and orf14 [3]. The S glycoprotein is embedded on the outer surface of the virus. By interacting with the angiotensin-converting enzyme 2 (ACE2) receptor, S protein trimers facilitate viral entry into host cells [4]. E protein plays an integral role in the production and maturation processes of the virus through interaction with the host cell membrane protein [5]. The M protein determines the shape of the E protein and contributes to the packaging of the viral RNA genome into a helical ribonucleocapsid in the process of virion formation [6]. The N protein, bound to the viral RNA genome, controls the viral replication cycle, the viral genome signaling, and the host cell reaction to the viral infection [5].

With the hope for a safe and effective strategy to tackle the COVID-19 problem, the interaction between drugs and structural proteins has been the subject of several therapeutic strategies [7-11]. Also, different vaccination pipelines have been developed mainly targeting the S protein [12, 13]. On the other hand, mutations of the structural proteins could increase the viral load, infectivity, and immune escape. For example, B.1.617.2 (Delta) and B.1.1.529 (Omicron) are more transmissible and resistant to neutralization due to mutations found in the structural proteins, especially the S protein [14-16]. Of note, the D614G, located in the S protein, is the most common variant of SARS-CoV-2, which increases infectivity [17]. These alterations may provide challenges in tackling the COVID-19 problem suggesting the importance of mutations in determining the quality of vaccinations and therapeutic strategies.

Our research aimed to determine the frequency of mutations in SARS-CoV-2 structural proteins in different American countries and also the entire world. We also investigated the evolutionary patterns of mutations based on their properties and ultimately, the mutations features will be compared between the Americas and worldwide data.

## Materials and Methods

### Sequence gathering

Mutation tracking was performed by gathering the SARS-CoV-2 AASs and comparison the data with Wuhan-2019 virus as reference sequence (access number: EPI_ISL_402124). Extraction of samples was conducted from GISAID database with the permission of Erasmus Medical Center [18-20]. All involved samples in the study were for human and samples with unknown time and location, different AAs length and non-specified AAs were kept out. Details of AA regions and the lengths of structural proteins were depicted in figure 1.

**Figure 1.**
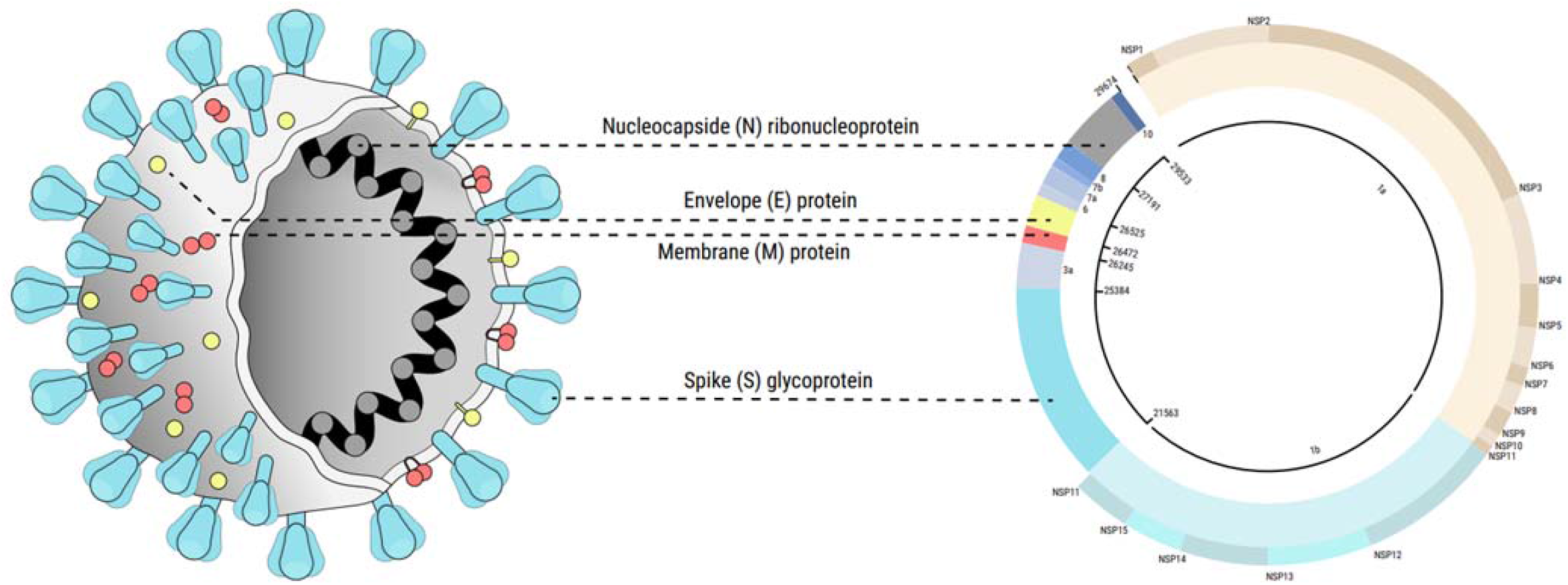
the lengths and regions of structural proteins in SARS-CoV-2. E, M, N and S protein are 75, 222, 419 an 1273 in length, respectively. The regions of 26245-26472, 26523-27191, 28274-29533 and 21563-25384 are the genomic positions of E, M, N and S, respectively.

### Sequences analyses

Sequences alignment and analysis was performed in the form of FASTA files with Python 3.8.0 software. As it mentioned above, Wuhan-2019 was utilized as reference sequence and mutation/s was introduced via detection of the difference/s between samples and this sequence. Optimization of data was completed by applying ‘Numpy’ and ‘Pandas’ libraries.

Mutation tracking was conducted using following algorithm:

~~~
> For refitem, seqitem in zip (refseq, seq)
If (refitem! =seqitem)
Report a new mutant
~~~

Because of the equal lengths among all sequences, the following algorithm used ‘Refseq,’ and ‘seq’ refer to reference sequence and sample sequence, respectively. Ultimately, reports contain mutations, locations and subset AA and the figures were drawn using R 4.2.0.

### Data normalization and statistical analysis

In order to compare the data with more accuracy, normalization was conducted using R 4.0.3 and Microsoft Power BI. Therefore, for each country, the number of mutations was divided by the number of attributed sequences that were comparable in equal proportions.

## Results

### Results related to the quantity of mutations

Investigating the GISAID database led to extracting 9670364 qualified samples from North America and 639746 eligible samples of AASs from South America. Bedsides, 26090908 AASs sequences belonging to structural proteins of SARS-CoV-2 were extracted globally. Samples of North America included 427550 samples for S AASs, 3597547 samples for E AASs, 3182356 samples for M AASs and 2462911 samples for N AASs. In comparison, there were 64546, 217344, 202987 and 153869 samples from South America attributed to S, E, M and N AASs, respectively. Additionally, worldwide samples included 950459 samples for S AASs, 9914529 samples for E AASs, 8860463 samples for M AASs and 6365457 samples for N AASs.

2.13% of S samples attributed to North America did not carry any mutation and 3.18%, 20.58%, 9.99% and 35.42% of them displayed one, two, three and more than three mutations, respectively (Figure 2). Considering the S sequences of South America demonstrated different situation, in which 80.52% of samples carried more than three mutations and only 0.89% of S AASs did not have any mutation. Furthermore, 4.82% of worldwide results displayed no mutation and 26.31%, 25.31%, 13.75% and 29.79% of such samples included one, two, three and more than three mutations, respectively. More than 99% of E AASs from both North America and South America displayed no mutation or maximum one mutation. Among samples of North America, 72.85% and 27.01% of them carried no mutation and one mutation, respectively and in almost similar results from South America, 76.93% and 22.93% of samples included no mutation and one mutation, successively. In addition, 67.72% and 32.09% of global E samples demonstrated no mutation and one mutation, respectively.

**Figure 2.**
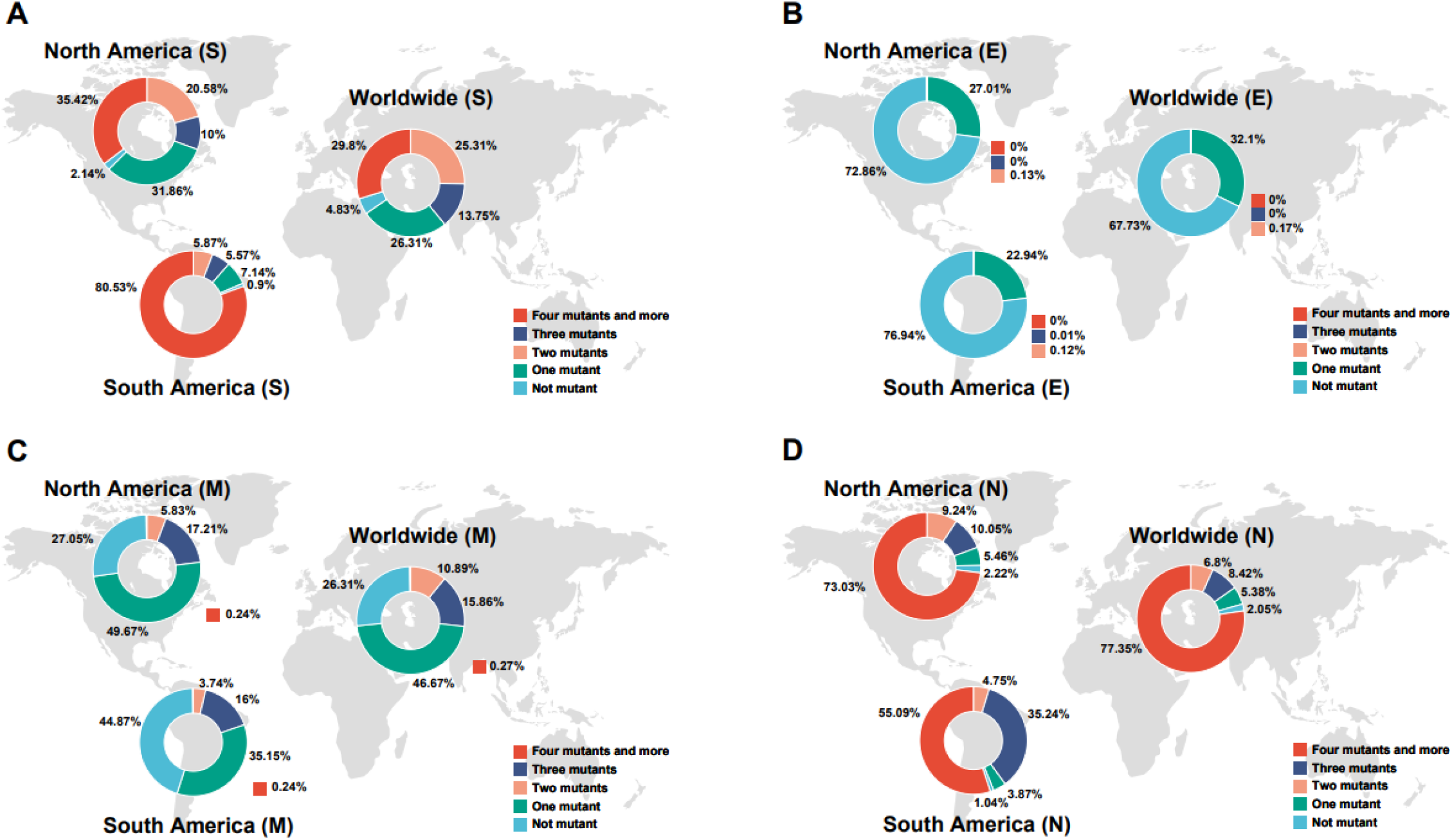
Pie chart plot of the number of mutations in S, E, M, N proteins of SARS-COV-2 as of April 2022. Results belonging to North America, South America and Worldwide samples were depicted in the sections of A, B and C, respectively.

Studying of M AASs demonstrated not incidence of mutations in 27.05%, 44.88% and 26.30% of samples belonging to North America, South America and entire world. Among sequences of North America, incidence rate of one, two, three and more than three mutations was concluded as 49.67%, 5.82%, 17.21% and 0.23%, successively. These former parameters for South America were observed as 35.14%, 3.74%, 16% and 0.23%, respectively. Moreover, analyzing the N AASs displayed no incidence of mutation in 2.21% of samples from North America and 5.46% and 73.03% of such samples included one and more than three mutations, respectively. Also 1.04% of samples belonging to South America did not display any mutation and almost 91% of such samples included three and more than three mutations, in which 35.24% had three mutations and 55.09% of them showed more than three mutations.

Results also were utilized in drawing a heat map for each of structural AASs to finding the regions with high predisposition toward mutation incidence. Although the region of 508 to 635 was concluded as the region with highest mutations frequency among S AASs from North America and also entire world, the most mutations relative to the total AASs among South America was occurred in the region of 1 to 127 (Figure 3). On the other hand, this different result was not observed in E samples, i.e. the region with highest mutations frequency was identical in all of them (7 to 14). Conversely, analyzing the data from North America and Worldwide samples introduced the region of 66 to 88 among M AASs as region with highest frequency rate. But highest frequency of mutations was occurred in the region of 1 to 22 in South America samples. On top of that, the region of 164 to 205 was concluded as region carrying highest frequency of mutation incidence in N samples from North America, South America and worldwide sources.

**Figure 3.**
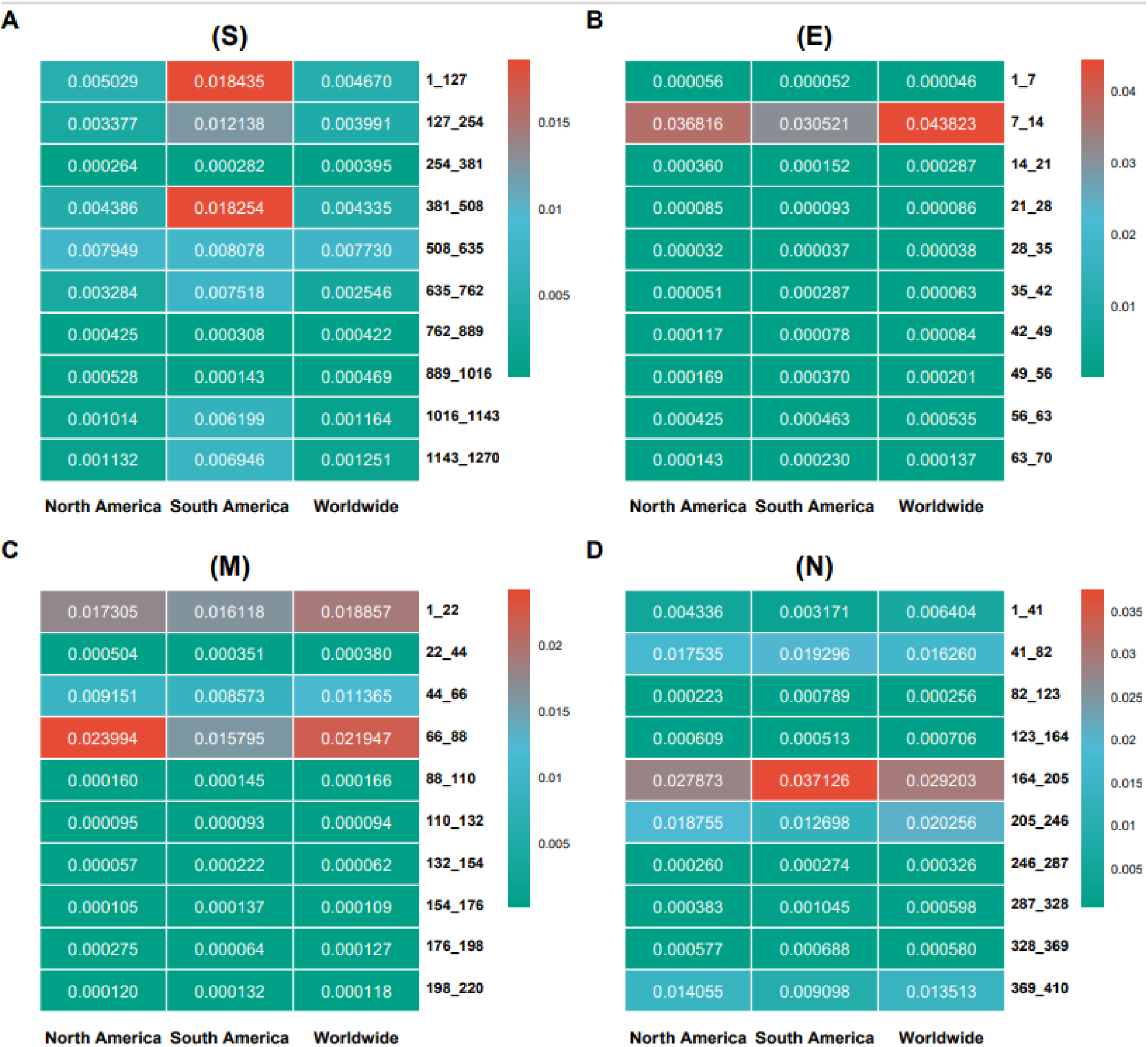
Heat map of hotspot regions in S (A), E (B), M (C) and N (D) AASs of SARS-COV-2 up to April 2022 among North America, South America and Worldwide. The heat map indicates the rate of mutation per 100 AAs.

### Discovered mutations among structural proteins

D614G was found as the most frequent mutation among S AASs in both North America and South America with almost similar frequency (0.9837 in North America and 0.9957 in South America) (Figure 4). But arrangement of subsequent mutations was different between sequences of these two continents. E484K (0.1357 frequency) and L18F (0.1204 frequency) were detected as second and third frequent mutations in S AASs from North America. On the other hand, V1176F with 0.8466 frequency and E484K with 0.8118 frequency were introduced as second and third frequent mutations in S AASs belonging to South America. Worldwide data demonstrated that D614G (0.9764 frequency), E484K (0.1406 frequency) and L18F (0.1623 frequency) were top three frequent mutations, globally. Furthermore, T9I (0.2574 frequency), P71L/S (0.0016/0.0008 frequencies) and L21F/V/I (0.0013/0.0006/0.0001 frequencies) were first to third frequent mutations in E proteins attributed to North America. Additionally, first and second frequent mutations among such proteins of South America were similar to mutations of North America, but third frequent mutation was observed as V58F with 0.0015 frequency. T9I with 0.3064 frequency, P71L with 0.0046 frequency and V62F with 0.0022 frequency were observed as first to third frequent mutations entire world.

**Figure4.**
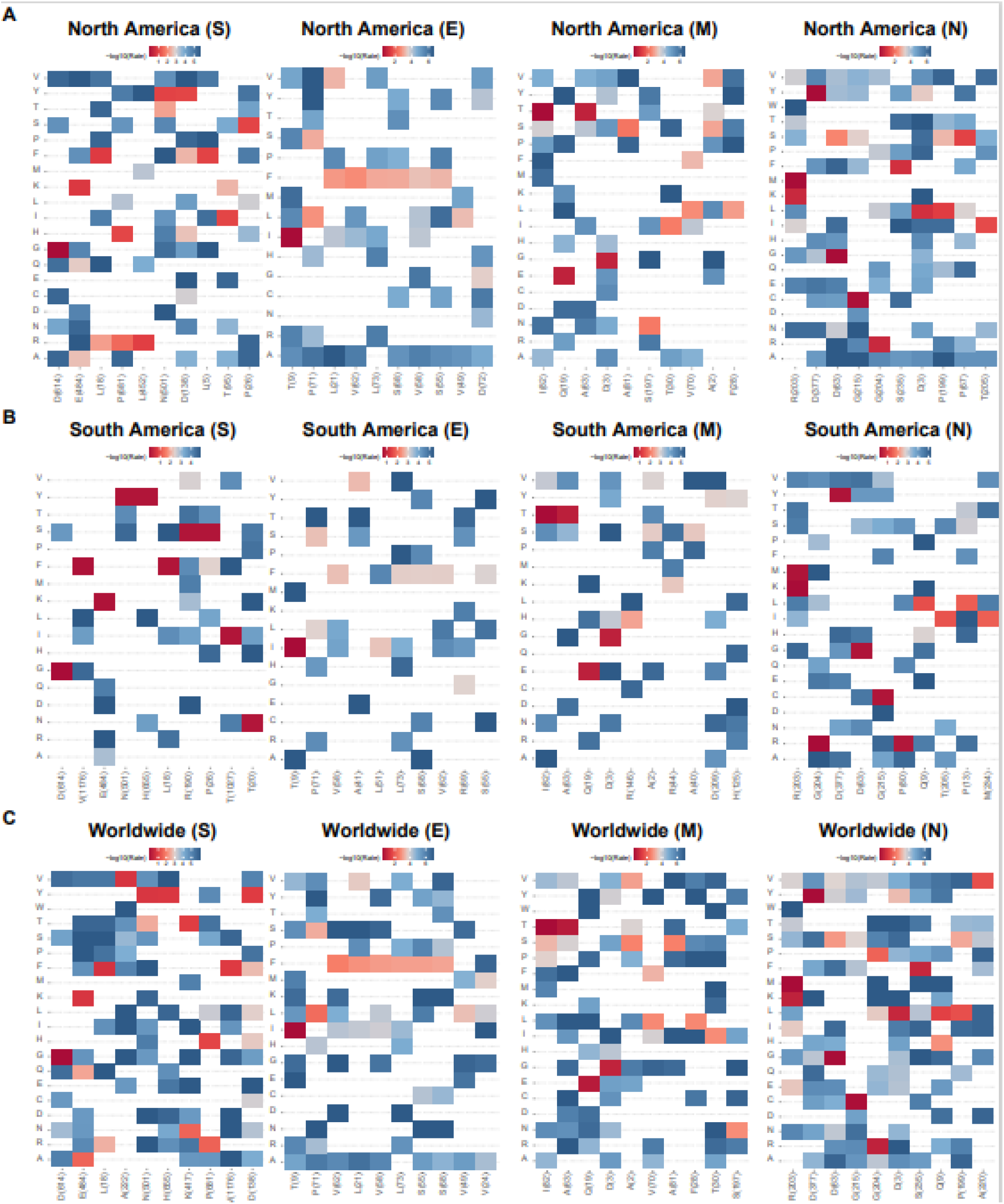
Top 10 mutations in S, E, M, N of SARS-COV-2 with the highest frequency In (A) North America, (B) South America and (C) Worldwide. The position of altered amino acids and substituted ones is shown differently base on the frequency percentage of substituted AA. The mutation frequency was estimated for each of them b normalizing the number of genomes carrying a given mutation in a desired geographic area.

Interestingly, although first three frequent mutations of M AASs from both North America and South America were similar in types of mutations, their arrangements are noticeably different. I82T (0.5121 frequency in samples from North America and 0.3416 Frequency in samples from South America) was the most frequent M mutation in both continents but Q19E (0.1990 frequency) and A63T (0.1986 frequency) were second and third frequent mutations in M AASs from North America. In comparison, A63T with 0.1869 frequency and Q19E (0.1777 frequency) were found as second and third frequent mutations in South America; which had identical arrangement with top three frequent mutations globally. Also, considering the mutation frequencies among N AASs were indicated that R203M/K (0.6457/0.1689 frequencies), D377Y (0.6562 frequency) and D63G (0.6404 frequency) were first to third frequent mutations in North America. Further, R203K/M (0.5161/ 0.4128 frequencies), G204R (0.5147 frequency) and D377Y (0.4178 frequency) were concluded as first three frequent mutations in South America. Ultimately, R203M/K (0.6282/0.2359 frequencies), D377Y (0.6329 frequency) and D63G (0.6202 frequency) were showed as frequent mutations entire world with similar arrangements to North America. Additional data about the Americas have been listed in supplementary files S-FREQ-AMR, E-FREQ-AMR, M-FREQ-AMR and N-FREQ-AMR and data belonging to the global samples have been listed in supplementary files S-FREQ-WRD, E-FREQ-WRD, M-FREQ-WRD and N-FREQ-WRD.

### Trends for emergence and distribution of structural mutations

Timeline analyses of mutations displayed similar pattern of emergence and distribution trend of D614G among both continents and also entire world (Figure 5). D614G frequencies gained to increase from February 2020 and up to now saved its maximum frequency in almost fluctuated trend. What’s more, from November 2021, T9I mutation gained to increase in both continents and received its maximum frequency in close to January 2022. Evolutionary patterns of M AASs showed that in January 2022, when I82T mutation was in next to minimum frequency, Q19E and A63T mutations had noticeably higher frequency than I82T and gained to increase in a growing trend started from November 2021. The similar trend was resulted in timeline of N AASs, i.e. when D377Y and D63G started decreasing trends of frequencies from December 2021, G204R gained to noticeable increase in both continents. Supplementary files S-EVL-AMR, E-EVL-AMR, M-EVL-AMR, N-EVL-AMR, S-EVL-WRD, E-EVL-WRD, M-EVL-WRD, N-EVL-WRD include supplemental data about the evolutionary patterns of mutations.

**Figure5.**
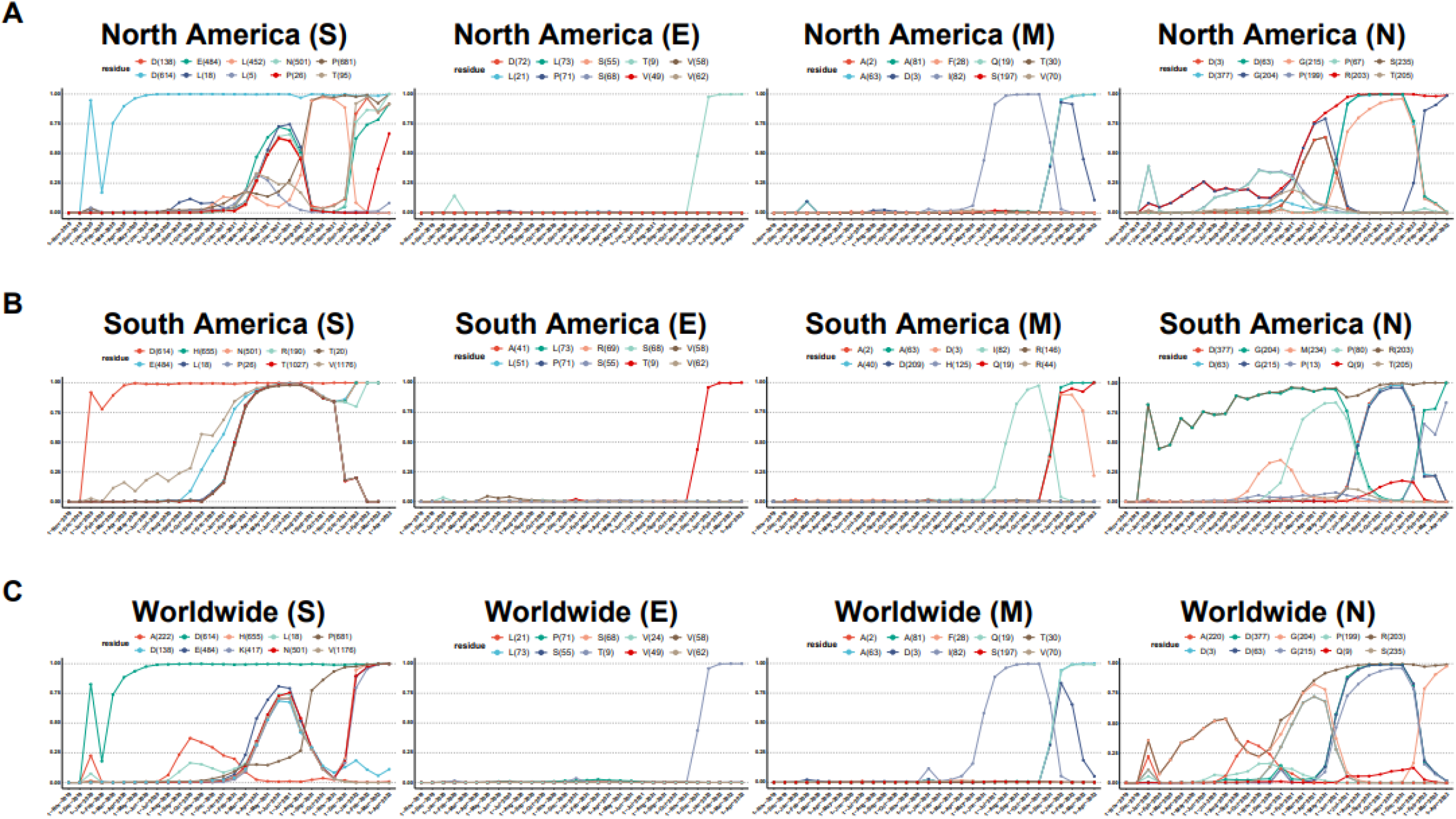
time evolution trajectories of top ten high-rate mutations of S, E, M, N of SARS-COV-2 in different geographic areas including (A) North America, (B) South America, (C) Worldwide. According to the month of sample collection, data is computed as the number of AASs having a given mutation over the total number of AASs according to the month of sample collection.

## Discussion

Keeping an eye on the evolution of SARS-CoV-2 and monitoring emerging mutations is essential for developing new diagnostic tests, vaccines, and treatments against COVID-19 in this ongoing pandemic. More than 90% of samples in each group had at least one mutation in the S or the N proteins in our study. On the other hand, the E and the M proteins had the least mutation rate. This is consistent with the data from Mutation Tracker [21]. According to Sato et al., the structural proteins had ∼99% conservation until 2020, with the S protein having the highest proportion of AA changes along its sequence [22].

Substituting aspartic acid (D) to glycine (G) in residue 614 of the S protein has been one of the most frequent mutations in the viral genome. Evidence shows that the D614G facilitates cell entry, replication, infection, and fitness [23, 24]. D614G enhances binding to ACE2 [24, 25]. The aspartic acid in position 614 resides on the surface of the S protein. A hydrogen bond could form between the side chains of this aspartic acid and that of its neighboring protomer, linking residues of the S1 unit of one protomer with residues of the S2 unit of the neighboring protomer. Substitution of D614 with glycine would disrupt this side-chain hydrogen bond [26], resulting in a more stable structure with more S protein incorporation into the pseudovirion [27]. We found that T9I is now the most frequent among the E protein mutations, while in a previous study, S68F was recognized as the most common mutation in the E protein [22]. The prevalence of this mutation in Omicron strain has been determined as ∼100% [28, 29]. The 9th position of E protein resides at the top of the transmembrane domain [30]. How the substitution of the hydrophilic threonine (T) to the hydrophobic isoleucine (I) affects the function of the virus is not precisely understood. However, it is suggested that this substitution could probably strengthen the interaction of the virion with membrane lipids because of isoleucine’s hydrophobicity [31].

Although the M protein has been mainly conserved throughout the pandemic till February 2021, it underwent rapidly increasing mutations. We observed that I82T mutation is the most prevalent in North and South America, with 0.5121 and 0.3416 frequencies, respectively. This is similar to previous studies indicating the I82T mutation as one of the most frequent mutations in the M protein in the USA, with a 116-fold increase from October 2020 to February 2021 [32]. Suratekar et al. also reported I82T as one of the most prevalent mutations in the Delta variant in a systematic analysis of mutations from nearly 2 million SARS-CoV-2 genomes from 176 countries [33]. However, till 2020, D3G was reported as the most common mutation in this protein [22]. The change of a hydrophobic amino acid (Isoleucine) to a more polar one (Threonine) has not been shown to alter the structure of the M protein; however, the similarity of the M protein structure to glucose transporters suggests that this mutation may contribute to glucose uptake during viral replication [32].

Consistent with our analysis, alteration of the residue 203 in the N protein has been reported as the most frequent mutation in this protein. Our analysis consistently showed that substitution mutation in the region of 203 was the most common change in the N protein in North America, South America, and worldwide. Mutations in this residue have been seen in Alpha, Delta, and Omicron lineages [34, 35], increasing the SARS-CoV-2’s infectivity and immune escape ability [34-37]. In an experiment with live a Coronavirus model, Delta variants with R203M could deposit 10 times more mRNA into host cells than wild-type viruses and generate 51 times more infectious viruses [37]. Also, another study showed that the R203M Delta variant had increased viral titer in patients [35]. This may be because R203 mutation increases the virion assembly by increasing the condensation of N-protein with viral genomic RNA [38].

Our analysis showed that the mutational hotspot regions in the S protein were positions 508 to 635 in North America and the whole world, the majority of which constitute the S1 region of the protein. In contrast, in South America, the N terminal domain (NTR), receptor-binding motif RBM), and receptor-binding domain (RBD) were the locations with the most prominent mutations (1-127, 381-508). Consistent with this, Harvey et al. showed that residues at positions 18, 222, and 614 had the highest variation frequencies [39]. The RBD is essential as it is the region that binds to ACE2 and is targeted by vaccination platforms. The efficacy of vaccines against the Omicron variant was heavily attenuated, providing a significant concern for protection against the disease [40]. One of the responsible mutations for this evasion is R346K [41]. Surprisingly, the frequency of this mutation has bluntly decreased in the recent months in the North America, according to our data (from 0.79 to 0.16 within two months). Also, via an artificial intelligence (AI) model, researchers have found that K417N, E484A, and Y505H in the S protein could cause the most significant disruption of known antibodies to the S protein [42]. The frequency of these mutations has been rising until April 2022, but, in fact, we found that E484K, K471T, and Y505J are the most frequent amino acid changes for the 484, 417, and 505 positions of the S protein. This information may help provide a more precise prediction on how the functional mutations affect vaccination immunity.

A limitation of this study was not considering the nucleotide sequences of the virus and only focusing on the amino acid sequences, neglecting the aspects such as codon bias [43]. Also, the sample sizes in the dataset are biased towards Europe and North America, suggesting that some functionally significant variants may be overlooked. Overall, COVID-19 vaccination and treatment success is determined by the frequency of functional mutations and the rate at which they appear. Faster identification of mutations could speed up the adopted approaches for combating COVID-19.

## Supporting information

E-EVL-WRD

E-FRQ-WRD

E-EVL-AMR

E-FRQ-AMR

M-EVL-AMR

M-FRQ-AMR

M-EVL-WRD

M-FRQ-WRD

N-EVL-AMR

N-FRQ-AMR

N-EVL-WRD

N-FRQ-WRD

S-EVL-AMR

S-FRQ-AMR

S-EVL-WRD

S-FRQ-WRD

## Acknowledgments

The authors thank all of the researchers who have shared genome data openly via the Global Initiative on Sharing All Influenza Data (GISAID).

## Additional information

### Supplementary Information

Supplemental data associated with the current study have been gathered.

